# Statistical classification of dynamic bacterial growth with sub-inhibitory concentrations of nanoparticles and its implications for disease treatment

**DOI:** 10.1101/2020.07.19.210930

**Authors:** A-Andrew D Jones, David Medina-Cruz, Na Yoon(Julie) Kim, Gujie Mi, Caterina Bartomeu-Garcia, Lorena Baranda-Pellejero, Nicole Bassous, Thomas J. Webster

## Abstract

Nanoparticles are promising alternatives to antibiotics since nanoparticles are easy to manufacture, non-toxic, and do not promote resistance. Nanoparticles act via physical disruption of the bacterial membrane and/or the generation of high concentrations of reactive-oxygen species locally. Potential for physical disruption of the bacterial membrane may be quantified by free energy methods, such as the extended Derjuan-Landau-Verwey-Overbeek theory, which predicts the initial surface-material interactions. The generation of reactive-oxygen species may be quantified using enthalpies of formation to predict minimum inhibitory concentrations. Neither of these two quantitative structure-activity values describes the dynamic, *in situ* behavioral changes in the bacteria’s struggle to survive. In this paper, borrowing parameters from logistic, oscillatory, and diauxic growth models, we use principal component analysis and agglomerative hierarchical clustering to classify survival modes across nanoparticle types and concentrations. We compare the growth parameters of 170 experimental interactions between nanoparticles and bacteria. The bacteria studied include *Escherichia coli*, *Staphylococcus aureus*, Methicillin-Resistant *Staphylococcus aureus*, *Staphylococcus epidermidis*, *Pseudomonas aeruginosa*, and *Helicobacter pylori*, and were tested across multiple concentrations of liposomal drug delivery systems, amphiphilic peptide, and silver and selenium nanoparticles. Clustering reveals specific pairs of bacteria and nanoparticles where the nanoparticle induced growth dynamics could potentially spread the infection through the development of resistance and tolerance. This rapid screening also shows that bacteria generated nanoparticles do not induce growth modes indicative of the development of resistance. This methodology can be used to rapidly screen for novel therapeutics that do not induce resistance before using more robust intracellular content screening. This methodology can also be used as a quality check on batch manufactured nanoparticles.

## Introduction

The post-antibiotic world is creating an economic and medical crisis with over 2 million hospital infections and 99,000 deaths, at a cost of $21 to $34 billion dollars in the US alone.^1,2^ Over 70% of hospital acquired infections are antibiotic resistant, some multi-drug resistant. Furthermore, after the introduction of a new antibiotic, it does not take long for bacteria to acquire or develop resistance.^3,4^ The discovery of new antibiotics is a slow process. This is, in part due, to a lack of incentives, the exhaustion of naturally derived antibiotics, and the biocompatibility of synthetic antibiotics.^4^ Changes in how we prescribe antibiotics, an increase in hygiene, sanitation and food safety practices may attenuate the crisis, however longer lifespans and a higher number of patients will increase the total risk of infection.^5^

Nanoparticles (NPs) are a relatively recent weapon in the fight against infection, with over 50 nanoscale drugs having been FDA approved already and many commercialized in the last 30 years.^6–9^ NPs may be designed to be, simultaneously, nontoxic, able to stay in the circulation and be cleared from the body, by modifying their size, shape, and charge.^9^ This has resulted in bare and functionalized polymeric drug carriers that transport existing antibacterial agents at lower concentrations and specifically target bacteria, which leads to a greater treatment efficacy and fewer side-effects than when the drug is administered alone.^6,10^ These carriers have lengthened the effective time existing antibiotics can be used, but have not fundamentally changed the antibiotic landscape.

Metallic and metal-oxide NPs, amphiphilic peptides, and nanotubes have been proposed in an effort to fundamentally change the antibiotic landscape.^6^ Because these NPs can be rapidly realized from rational design relative to antibiotic development, most modeling efforts have focused on them.^9,11^ However, many of the calculated and experimental parameters, such as clearance rates, have not translated well.^9^ Furthermore, variability in size, shape and charge from manufacturing processes and between *in vitro*, *in vivo*, and *in situ* environments resulted in conflicting reports, as similarly composed NPs have been reported to be both toxic and nontoxic to human cells.^9,12^ Furthermore, quality control of NP production is challenging without intentional development ecosystems,^8^ necessitating fast screening tools, such as those used to check for antibiotic contamination.^13,14^

NP usefulness as antibiotics and their potential toxicity can be quickly screened if their mechanism of action is known. NPs are proposed to act through the release and/or production of chemical species, mechanical interference like poration or binding, and enhanced permeation leading to further mechanical or chemical cell damage^15,16^. Unlike many antibiotics, they will act even when the cell is not dividing. The enthalpy of formation of chemical species can predict antibacterial activity for metal oxides.^15^ The enthalpy of formation does not account for protein coronas or other facets of the *in vivo* environment, nor describes innate or induced resistance. The extended Derjuan-Landau-Verwey-Overbeek (XDLVO) theory can predict nanoparticle agglomeration, which is one method of antibiotic resistance.^17,18^ XDLVO has also been used to predict adhesion of cells and proteins to surfaces.^19^ XDLVO includes surface roughness, acid-base chemistry, through contact angle measurements and zeta-potential of bacteria, nanoparticles and the growth media. However, surface energy does not describe the release of chemical species and can be cumbersome as material-free energies cannot be tabulated like enthalpies. A robust understanding of membrane cell interaction has been built from molecular dynamic simulations of nanomaterial-biological interaction, however these simulations require extensive time and knowledge of the material.^20^

To overcome these limitations, statistical tools have been proposed to supplement the wide variety of uncharacterized nanoparticle/bacteria interactions.^21^ For example, multivariate linear regression and linear discriminant analysis were used to distinguish the role of five material parameters on cell cytotoxicity.^22^ Counter propagation artificial neural networks were recently used to study the cytotoxicity of 72 metal oxide nanoparticles against *E. coli* in the context of developing a framework for environmental and health regulation.^23^

Here, we follow methods similar to Sayes et al.,^24^ combining principal component analysis and a clustering technique, specifically hierarchical agglomerative clustering, to differentiate the behaviors and classes of nanoparticle-bacteria interactions. Unlike many existing analyses, our dataset includes many different bacterial species and a combination of metallic and non-metallic nanoparticles. Furthermore, we apply the analysis to dynamic growth measurements similar to ASTM guide E2149-13a, instead of to a minimum inhibitory concentration, acknowledging the importance of time to both inhibition and resistance.^2,25^ Furthermore, while time series measurements may not reveal a mechanism, they can narrow the window required for future mechanism searches. We explore five parameters derived from three distinct growth models: Gompertz’, or Huang’s, model of logistic growth; an underdamped model of oscillatory growth; and Liquori’s model of diauxic or two-phase growth. We use various concentrations of commercial and green-synthesized metallic nanoparticles, amine-capped nanoparticles, bare and functionalized polymeric capsules containing nanoparticles and antibiotics. Using principal component analysis, we show that species and membrane structure cannot explain all the variance in nanoparticle-bacteria interactions, and that hierarchical agglomerative clustering reveals correlation in the behavior of unrelated NP-bacteria pairs.

## Methods

In order to classify nanoparticle-bacteria interactions for their potential to induce resistance or to promote bacteria proliferation, we analyzed 170 published and non-published interactions studied in our lab. The duration of each interaction was 24 h in a plate reader. The growth curves were fit to two growth models that describe logistic growth, and to growth models that describe diauxic and oscillatory growth. The quantitative parameters from these models were then used in the principal component analysis, and the transformed quantities classified using hierarchical agglomerative clustering.

### Growth Model Selection

Bacteria exposed to sub-inhibitory concentrations of nanoparticles can exhibit three growth behaviors, as shown in Figure 1. Logistic growth is seen under standard conditions and has been modeled with varying degrees of accuracy.^26^ Diauxic, or two-phase, growth occurs when bacteria are able to use a secondary compound or have overcome an inhibitory compound.^27^ It has also been identified as a sign of bacterial tolerance.^28^ Oscillatory growth is predominately seen in predator/prey situations, although we also observed this behavior in some of our nanoparticle-bacteria interactions and it has been implicated in bacteria free-riding on other bacteria’s resistance.^29^

**Figure 1.**
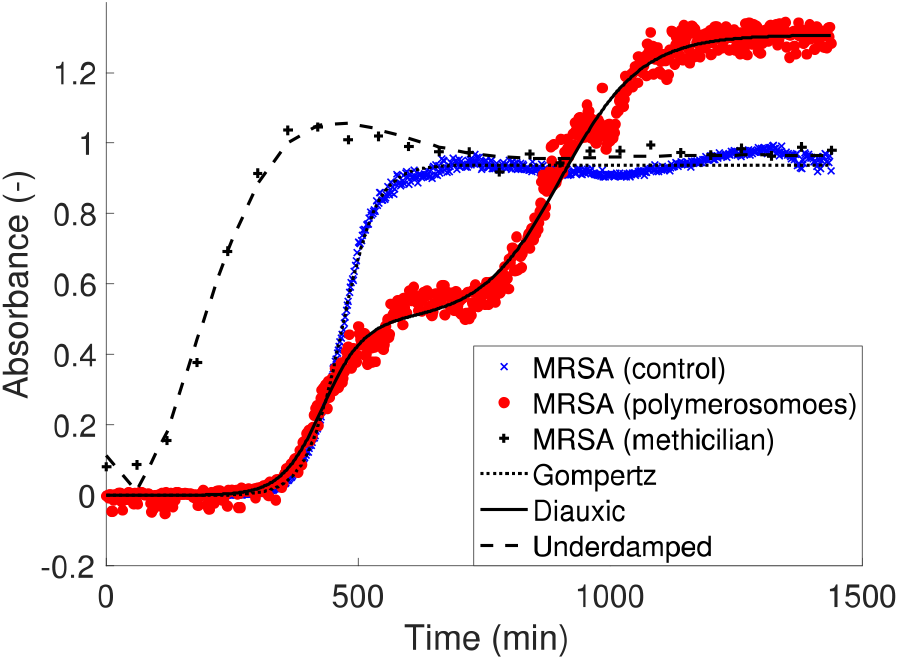
MRSA showing Gompertz, Diauxic, and underdamped oscillatory growth behavior when subjected to two different nanoparticles. Lag time, growth rate, carrying capacity, overshoot, and diauxic growth partition coefficient parameters can be extracted by fitting.

The modified Gompertz growth model,^26,30^ Table 1.1, is known to accurately fit physical parameters of growth rate, *μ*, and carrying capacity, *A*, where *c*_0_ is the initial concentration. However, while it has moderate success at modeling the lag-time, *λ*, its first derivative is never null and therefore cannot accurately predict the lag-time.

**Table 1.** Models used for fitting data and the free parameters extracted from each model. The first two equations model the same logistic growth behavior with varying degrees of reported accuracy, the latter two equations model two-phase and oscillatory growth respectively.

| # | Source | Equation | Free Parameters |
| --- | --- | --- | --- |
| 1. | Gompertz<sup>26,30</sup> | $c_0 + A \exp\left(-\exp\left(-\frac{\mu e}{A}(t - \lambda) + 1\right)\right)$ | $c_0, A, \lambda, \mu$ |
| 2. | Huang<sup>26,31</sup> | $c_0 + A - \log\left(e^{c_0} + (e^A - e^{c_0})e^{-\mu\left(t + \frac{1}{4}\log\frac{1+e^{-4(t-\lambda)}}{1+e^{4\lambda}}\right)}\right)$ | $c_0, A, \lambda, \mu$ |
| 3. | Liquori<sup>27</sup> | $c_0 + Ax \frac{1 - \exp\left(-\frac{t}{t_{\alpha_1}}\right)}{1 - \exp\left(-\frac{t}{t_{\alpha_1}}\right) + \exp\left(-\frac{t}{t_{\alpha_2}}\right)} + A(1-x) \frac{1 - \exp\left(-\frac{t}{t_{\beta_1}}\right)}{1 - \exp\left(-\frac{t}{t_{\beta_1}}\right) + \exp\left(-\frac{t}{t_{\beta_2}}\right)}$ | $c_0, A, t_{\alpha_1}, t_{\alpha_2}, t_{\beta_1}, t_{\beta_2}, \mu$ |
| 4. | Oscillatory | $c_0 + A - \frac{\mu}{\omega} e^{-\zeta\omega(t-\lambda')} \left( 2\zeta \cos\left(\omega\sqrt{1-\zeta^2}(t-\lambda')\right) + \frac{2\zeta^2-1}{\sqrt{1-\zeta^2}} \sin\left(\omega\sqrt{1-\zeta^2}(t-\lambda')\right) \right)$ | $c_0, A, \lambda, \mu, \zeta, \omega$<br>$\lambda' = \lambda + \frac{A}{\mu} - \frac{2\zeta}{\omega}$ |

Huang’s growth model^26,31^, Table 1.2, is also a three-parameter model, like Gompertz’ model, but can model the lag-time. It has been shown it can predict parameters equivalent to the ones predicted by the more complex Baranyi’s growth model.^28^

**Table 2.** Descriptions of clusters predicted by hierarchical agglomerative clustering applied to the time-series growth parameters of bacteria interacting with nanoparticles.

| Cluster | Experimental ID | Predominate growth mode |
| --- | --- | --- |
| 1 | 20 | Diauxic |
| 2 | 146,149 | Diauxic |
| 3 | 54 | Logistic |
| 4 | 46,107 | Oscillatory |
| 5 | 100,103,105,106,122,123,124,125 | Logistic |
| 6 | 4,25,90,109,110,112,115 | Logistic |
| 7 | 2,3,6,24,29,30,31,32,33,34,35,36,39,40,41,42,43,44,47,48,50,52,59,61,63,65,68,71,73,74,75,85,86,88,91,93,94,99,108,113,117,118,119,120,126,138,139,140,141,142,143,144,145,147,148,150, 151,152,158,159 | Logistic, Diauxic |
| 8 | 14,15,18,49,57,60,62,132 | Oscillatory |
| 9 | 10,13,17,19,101 | Oscillatory, Logistic |
| 10 | 1,5,7,11,12,16,22,23,28,37,38,45,56,58,64,66,67,69,70,72,76,77,78,79,80,81,82,83,84,87,92,95,96,97,98,102,104,111,114,116,121,127,128,129,130,131,133,134,135,136,137,154,155,156,157,160,161,162,163,164,165,166,167,168,169,170 | Logistic |

The diauxic, or two-phase growth, is less commonly used in the literature,^32^ and describes a secondary growth phase where bacteria are able to use a secondary nutrient source, or have overcome some inhibitory compound.^33,34^ We have observed such growth dynamics with bacteria in the presence of nanoparticles. Liquori’s diauxic growth model,^27^ Table 1.3, was chosen over more accurate dynamic models of diauxic growth^33–36^ because those other models require knowledge of substrate utilization and/or gene regulation. It is inappropriate to map the growth rates and lag-times of logistic growth onto diauxic growth as there are two phases where those parameters exist. Among the seven parameters in the model in Table 1.3, the partition coefficient, *x*, describes whether the growth is dominated by the first phase or by the second phase, if the following criteria is met 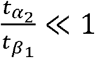 and. 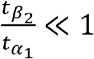.

Oscillatory growth describes the population overshooting, then oscillating about a carrying capacity. We derived biologically relevant parameters from the response of a second-order underdamped oscillator to a step-impulse, following the method of Zwietering et al.,^30^ as shown in Table 1.4. The additional parameter, *ω*, describes the frequency of oscillation about the carrying capacity. The damping ratio, *ζ*, is used to find the population overshoot, the population growing beyond sustainable limits, from controls theory as, 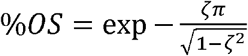

### Model Fitting

Models were fit using Trust-Region Reflective Least-Squared algorithm in MATLAB (Mathworks, Burlington, MA, USA). The lag-time, growth rate, and carrying capacity were constrained to physiological realistic, non-negative values. If possible, the parameters were initialized using results from a Levenberg-Marquardt fit of a logistic growth curve. Fits were accepted if *R*^2^ > 0.8. The Akaike Information Criteria was used to determine which of the models produced the best fit taking into account the number of free parameters. Here, we approximated the log maximum likelihood, 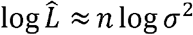, with the number of samples times the log of the variance.

### Experimental Conditions

In this analysis, we used existing published and unpublished data from our lab to examine the effects of nanoparticle exposure to seven different species of bacteria. All experiments were conducted in a 96-well plate plate-reader (SpectraMaxV R ParadigmV R Multi-Mode Detection Platform) for 24 hours. Methicillin-Resistant *Staphylococcus aureus* (MRSA) (ATCC 43300) were grown in tryptic soy broth at 37 ºC, exposed to liposomes containing methicillin, and functionalized with trans-activating transcriptional activator peptide. MRSA (ATCC 43300) were grown in tryptic soy broth at 37 ºC, exposed to temperature responsive polymersomes, poly-DL-lactic acid-(carboxyethyl) polyethylene glycol (PDLLA-PEG-COOH), encapsulating methicillin or silver nanoparticles and/or functionalized with 40 μM proline rich arginine. *E. coli* (ATCC 25922), *P. aeruginosa* (ATCC 27853), *S. epidermidis* (ATCC 35984), and *S. aureus* (ATCC 12600) were grown in tryptic soy broth at 37 ºC, exposed to amphiphilic peptides, as previously reported.^10^ *E. coli* and *S. aureus* were grown in LB broth at 37 ºC, exposed to selenium nanoparticles produced by exposing *E. coli*, *P. aeruginosa*, *S. aureus* and MRSA to selenium salts, as previously reported.^37^ *Helicobacter pylori* (NCTC 11637) were grown in tryptic soy broth at 21 ºC, exposed to selenium nanoparticles produced by exposing *H. pylori* to 1, 2, and 5 mM of Na_2_SeO_3_. *S. aureus*, MRSA, *E. coli*, MDR *E. coli* were grown in tryptic soy broth at 37 ºC, exposed to glutathione capped nanoparticles, liposomal cysteine capped silver nanoparticles, liposomal glutathione capped silver nanoparticles, and cysteine capped silver nanoparticles. Detailed parameter pairs are included in SI Table 1.

### Statistical Analysis

We applied two statistical data mining techniques, principal component analysis and hierarchical agglomerative clustering, to classify the behavior of the nanoparticle-bacteria interaction. Each observation had, at most, five parameters: lag-time, growth rate, carrying capacity, overshoot, and partition coefficient, which are the features used in our principal component analysis; the empty observations were filled in with the mean value obtained from the data set. We computed the relative lag-time, relative growth rate, and relative carrying capacity with respect to the control exposure, no nanoparticles, as (*ξ* - *ξ*)/*ξ*_0_. As our control exposure exhibited logistic growth for all species and media tested, it would be inappropriate to compute a relative overshoot and relative partition coefficient, however those values are measured relative to a logistic growth or single-phase growth, respectively.

Using principal component analysis, we discovered nanoparticle-bacteria pairs that warrant further study and potentially rank interactions. The principal component analysis is an approach used to maximize variance and show patterns between features of a data set. As our data set includes many different species and nanoparticle type, we do not expect it to produce a quantitative structure-activity relationship. However, because our data set is the largest studied to date, we hypothesized that it might reveal behaviors that might have been hidden among the other features. We used a centered standardized covariance matrix as there were multiple bacteria-nanoparticle interactions that produced relative lag-times that were orders of magnitude greater than the mean.

In order to determine which types of nanoparticle-bacteria interactions are unique, we used a clustering algorithm, Hierarchical Agglomerative Clustering. Hierarchical agglomerative clustering measures the similarity between independent nanoparticle-bacteria interactions. We used Euclidian distance over a centroid linkage to measure similarity.

## Results

Our methodology establishes a fast screening technique for classifying bacteria growth behavior in the presence of nanoparticles, using as inputs measurements of bacteria growth by optical density. These inputs are used in principal component analysis and hierarchical agglomerative clustering to separate out nanoparticle-bacteria interactions that exhibit tolerant, resistant, or persistent behavior. We explore explanations based on membrane and particle type. We use two models that describe logistic growth, Gompertz’ and Huang’s models; a model that describes diauxic or two-phase growth, Liquori’s model; and an oscillatory growth model. Using a best-fit approach, largest *R*^2^, Liquori’s growth model was selected as the model that best described a given behavior, except in cases of extreme oscillation. This is because there are seven free parameters instead of five free parameters of the oscillatory growth and three free parameters in Gompertz’ and Huang’s model. Using the Akaike Information Criteria,^31,38^ with the log maximum likelihood approximated as 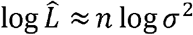, with the number of samples times the log of the variance, Liquori’s growth model still produces the best-fit as the parameter penalty is not large enough with respect to *n*, Figure 2a. Therefore, we assume a species can only be best-fit by Liquori’s growth model if 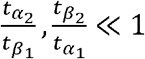 assuring a secondary plateau. Two-phase growth was exhibited by 52, or 30%, of the interactions spanning all the nanoparticle types except those generated by bacteria. If we exclude Liquori’s model from consideration, Figure 2b., Gompertz’ growth model describes 59% of the interactions. Huang’s model describes 16, or 9%, of the interactions, providing better estimates of the lag-time. Similarly to Liquori’s growth model, the penalty on the five parameters of the damped model is not large with respect to *n* as to discount its fits over the remaining 3 parameter models. However, if we include Liquori’s growth model with the constraint previously proposed, this over fitting is reduced.

**Figure 2.**
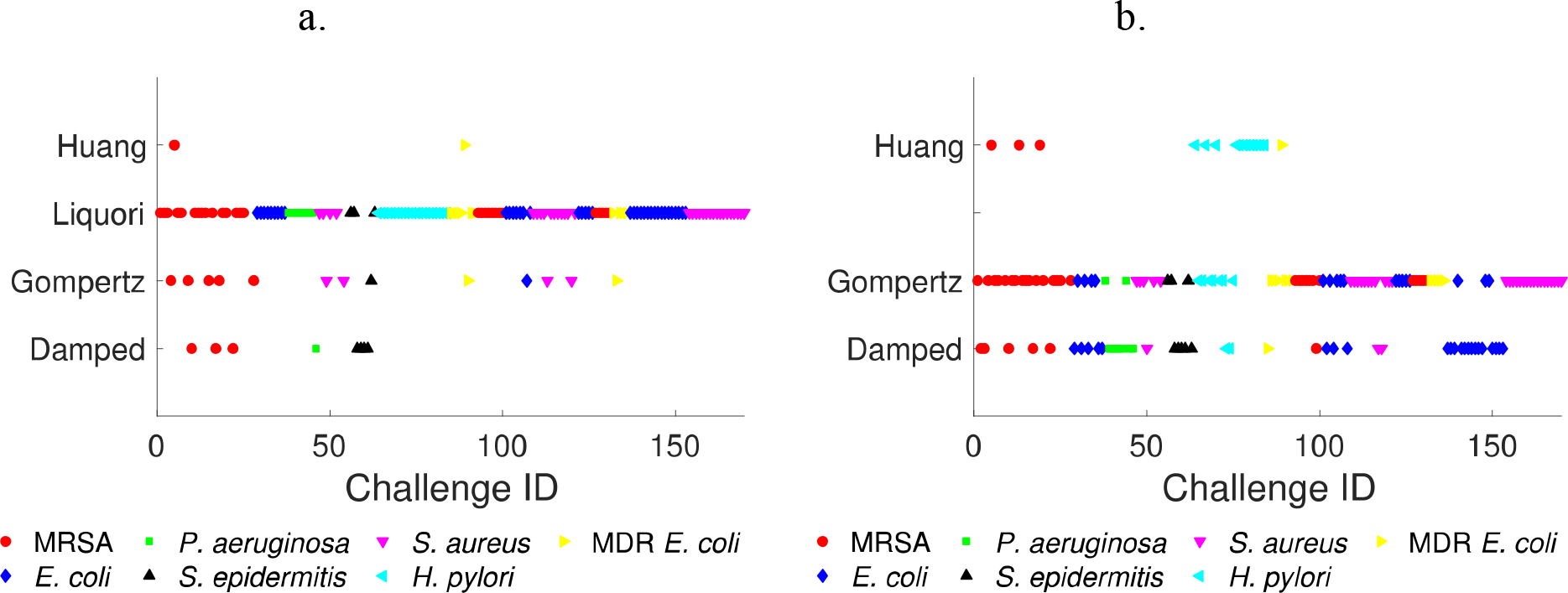
a. Minimum AIC of bacteria-nanoparticle interactions comparing all four growth models. b. Minimum AIC of bacteria-nanoparticle interactions excluding the Liquori’s growth model for two-phase growth.

From the model fits described above, we were able to derive relative growth rate, lag-time, and carrying capacity and report the percent overshoot and partition coefficient, Figure 3. The relative lag-time, (Figure 3 or SI Figure 1), is a measure of the time delay that the nanoparticles may induce if they posses antimicrobial properties. The high variance of the time delay is expected given that some of the concentrations of the nanoparticles are near the minimum inhibition concentration, while others do not possess antibacterial properties at all. We did find some of the nanoparticles induced shorter lag-times, which may be useful in combination therapies, as it has been found that higher metabolisms lead to greater susceptibility to antibiotics. The high variance in relative growth rate (SI Figure 2) and carrying capacity (SI Figure 3) are similarly understood as a function of near MIC concentrations. The percent overshoot (SI Figure 4) and partition coefficient (SI Figure 5) are induced behaviors so their relative quantities are meaningless. Furthermore, they are constrained to a range of [0,1] by definition.

**Figure 3.**
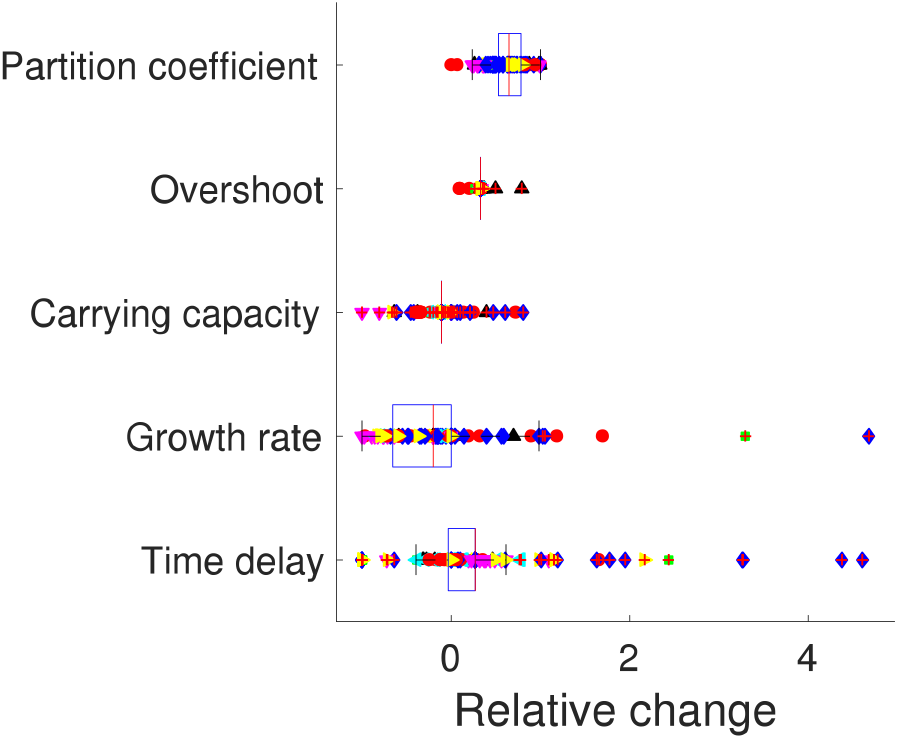
Box plot of the five parameters extracted from growth models of bacteria-nanoparticle interactions.

This work explores novel nanoparticles that do not have an easily quantifiable chemical composition, such as selenium nanoparticles created by bacteria^37^ or nanoparticles that require computationally expensive molecular dynamic simulations, such as amphiphilic peptides.^10^ This precludes the use of input parameters used for metallic nanoparticles, however, we still follow the methods of Sayes et al.,^24^ regarding the use of the principal component analysis, and also use clustering techniques. Our second objective was to develop a framework that will allow us to predict the development of resistance or tolerance, without using the minimum inhibitory concentration as an output. Instead, the input parameters to the model were taken from growth curves. We used a centered parametric matrix as the input for principal component analysis to account for the high variance in the data shown in Figure 3. As shown in the Pareto plot, Figure 4b, principal components 1 and 2 explain 50% of the variance with strong (> 30%) principal component 1 dependence. The control interactions cluster near the origin as should be expected. However, this breakdown also shows that these input parameters are not sufficient to explain the variance. Furthermore, principal component analysis is a form of factor analysis that does not produce quantitative explanations of the resulting factors. It is a data exploration tool that requires further inspection to produce quantitative results.

**Figure 4.**
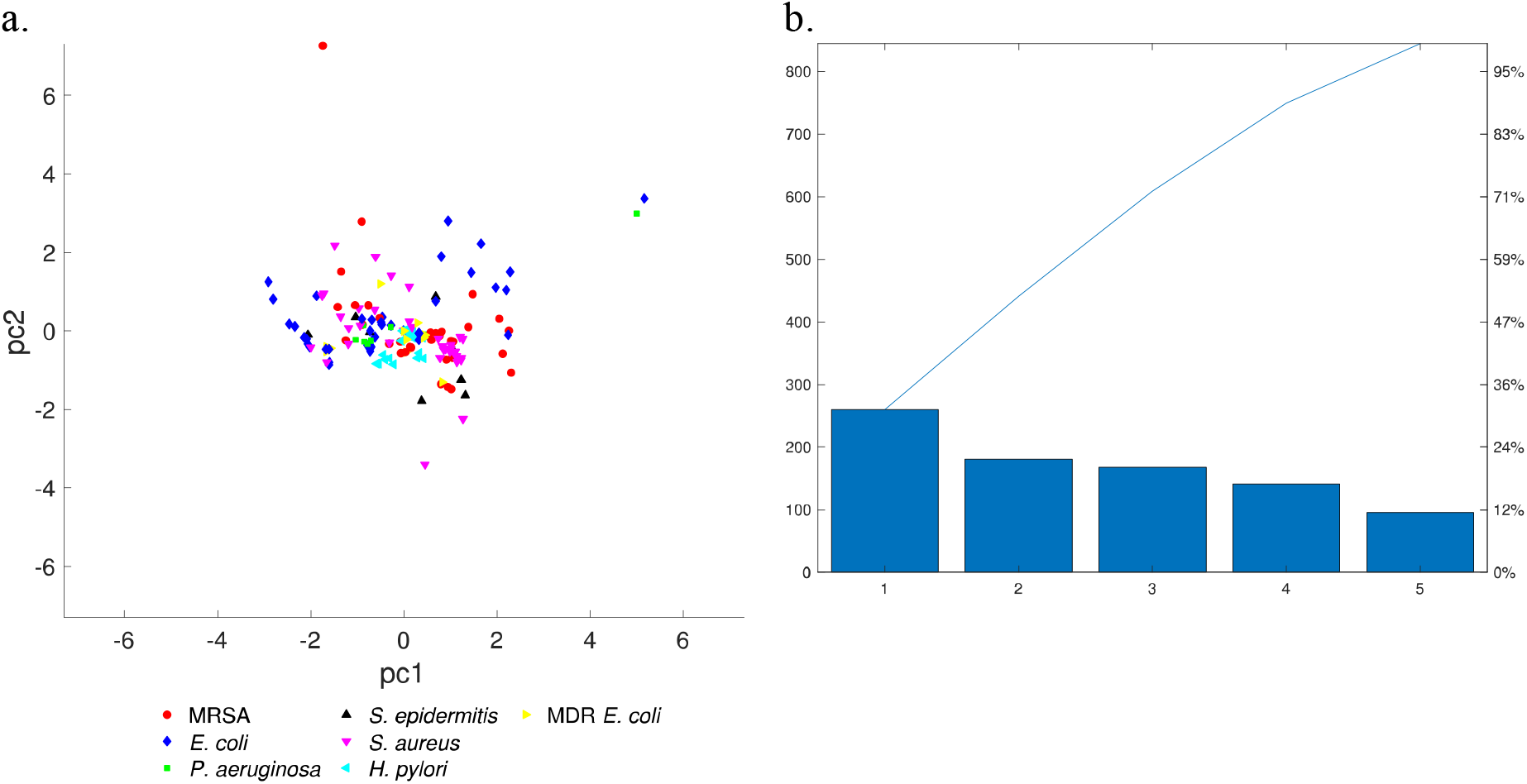
a. Principal component 1 versus principal component 2 plot highlighting bacterial species. b. Pareto plot of principal components shows pc1 and pc2 explain 50% of the variance of the data.

A clustering algorithm was used to further distinguish nanoparticle-bacteria interactions. Using hierarchical agglomerative clustering with a centroid linkage, we distinguish 10 unique clusters, Figure 5a. The clusters contain multiple species that do not correlate with the membrane structure described by Gram-staining, as shown in Figure 5b. The clusters are initially described by the predominate growth mode, as shown in Table 2.

**Figure 5.**
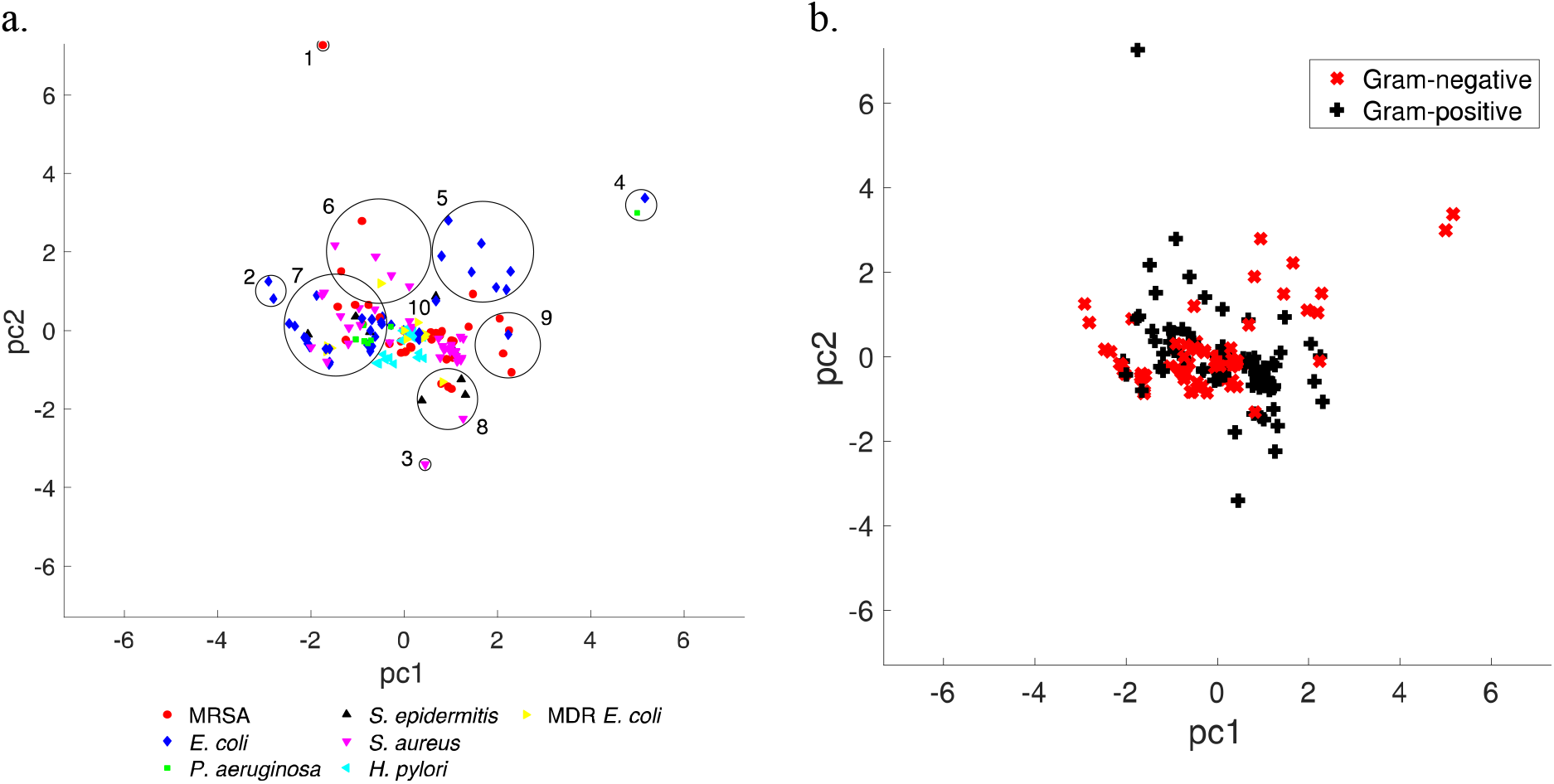
a. Hierarchical agglomerative clustering of the PCA transformed components using the centroid shows six distinct outliers surrounding a central cluster. The overlap in clusters 6 and 7 is strictly a function of the graphical representation. b. The membrane structure via Gram-stain overlaid onto the PCA shows weak correlation with the results from HAC.

Nanoparticle-bacteria interactions in clusters 4, 8, and 9 predominately exhibited oscillatory growth. The predominant species in these interactions is MRSA (47%) and the predominant nanoparticles are methicillin based nanoparticles (53%). Some of the interactions that visually appeared oscillatory were not classified by the fitting algorithm described above as oscillatory, for example, *E. coli* with cysteine capped silver nanoparticles (SI Table 1, EID 107), however, the additional parameters were sufficient to cluster them together. Other interactions have similarities that are hidden in the principal component analysis. The off-axis clustering is expected because this growth behavior is rare, but not desirable. Oscillatory growth has been found in the development of cooperative resistance of antibiotic treatment, when some parts of the population are resistant and others are “cheating.”^29^

The single-clusters, cluster 1 and cluster 3, are nearly inhibited interactions. MRSA exposed to a 3.3 μg/mL suspension of TAT coated liposomes (SI Table 1, EID 20) containing methicillin and resulting in diauxic growth exhibited a difference in absorbance of 0.2. *S. epidermidis* exposed to a 74.01 μg/mL suspension of amphiphilic peptides (SI Table 1, EID 54) exhibited a difference in absorbance of 0.06. In addition to the relative carrying capacity, a parameter expressing the absolute difference between initial and final cell density of colony forming units for future work could be used as a measure of inhibition, though such classification was not the intent of this study.

Cluster 5 predominately exhibits second growth phase dominate diauxic growth, with an average partition coefficient *x* = 0.69 ± 0.13. Cluster 6 exhibits delayed logistic growth, including a 400%-time delay of MRSA in the presence of 1.7 μg/mL of TAT coated liposomes containing methicillin (SI Table 1, EID 25) with only a 3% change in carrying capacity. The predominate strain in cluster 5 is *E. coli,* 87.5%, while in cluster 6 is *S. aureus*, 57.1%. Both clusters predominately consist of liposome nanoparticles. It has been found that tolerant bacteria exhibit this diauxic growth. ^28,39^ However, as our results come from drug carrying liposomal nanoparticles it may be possible to add sugars or other metabolic treatments to reduce or eliminate this behavior further extending the usefulness of the drug treatment. ^40,41^

The nanoparticle-bacteria interactions of selenium nanoparticles produced by bacteria are evenly split between clusters 7 and 10, though cluster 10 has more interactions in total (76 and 60 interactions, respectively). The additional interactions in cluster 10 are largely due to control species existing at the origin. Cluster 10 exhibits predominately logistic growth while cluster 7 splits between diauxic and logistic growth. As mentioned earlier, none of the bacteria produced nanoparticles induced diauxic growth.

## Discussion

Here we present categorization of nanoparticle-bacteria interactions using principal component analysis and hierarchical agglomerative clustering. Unlike molecular dynamic simulations or machine learning methods, clustering methods do not require a physical understanding as inputs however, they do not provide mechanistic explanations. As our data show, our method does provide a rapid reduction of a large data set with many complex interactions to a smaller data set that can be further studied with additional tools.

Understanding the nanoparticle-bacteria interactions on industrial and medical nanoparticles is important in consequential life-cycle analysis in order to balance indirect changes in multiple systems.^13,14^ A focus on time-series parameters, as opposed to inhibition, may support efforts to reduce the evolutionary selective pressure of future antibiotics.^3^ For example, our finding that bacteria produced nanoparticles did not induce secondary growth may be useful in industrial and medical applications for regenerative medicine^5^ even though we did not find significant antibacterial affects. Future work on categorizing bacteria-nanoparticle interactions from time-series extracted parameters may provide data enrichment for costly PCR monitoring of nanoparticle effects on mixed culture bacteria populations in nature, the human microbiome or wastewater activated sludge. In the same way that neural networks were used to propose recommendations for registration, evaluation, authorization, and restriction of chemicals legislation,^23^ we have met four out of the five principles specified by the OECD.^42^ Fast screening categorization can be used in quality control for properties such as size and surface charge, which correlate with the efficacy and persistence of the nanoparticles.^6,8^

Optical density measurements of bacterial growth time series are rapid and data rich, quantifying phenomena such as population overshoot or diauxic growth, ^32^ requiring minimal manual input as a fast screening method for nanoparticle-bacteria interaction. The data sparse quality of CFU counts hides information about intermediate phenomena, for example, treatment of hospital acquired strains of *P. aeruginosa* with silver NPs showed limited increasing growth at 0.156 and 1.25 μg/mL, and a sudden decrease at 5 μg/mL.^43^ Furthermore, this technique does not neglect agglomeration, surface charge, aqueous diameter, solubility, or protein coronas, which reduce the efficacy of nanoparticles *in vitro*. However, it is understood that there is not a one to one relationship between the optical density values and bacteria viability. Recently, dynamic light scattering was shown to be as accurate as plate counting to quantify viable cells, so it is foreseeable that DLS might be able to provide similar data with a one to one correlation with cell viability.^44^ Therefore, while the data quality herein may not be transferable, the parameters and models used to extract time-series behavior for fast-screening of resistance development and other undesirable behavior of antibacterial agents are likely transferable.

In this study, we did not use quantitative descriptors of the nanoparticles, such as hydrodynamic diameter or zeta-potential. In future studies, the inclusion of particle hydrodynamic diameter ratios to minimum bacteria radius, and zeta-potentials ratios may provide further explanation of bacteria-nanoparticle behavior.

## Conclusion

Principal component analysis and hierarchical agglomerative clustering were used to analyze data from over 100 experiments with bacteria exposed to nanoparticles in order to extract features and behaviors that are unique and warrant further study. The clusters did not map onto gram-staining. Instead, the clusters screened for certain bacteria nanoparticle interactions that exhibited oscillatory and diauxic growth, previously implicated in the development of drug resistance and tolerance. With 170 interactions, some bacteria-nanoparticle interactions that did not exhibit resistance or tolerance growth modes were clustered with those exhibiting resistance or tolerance, which warrants further study. We found that bacteria generated metallic nanoparticles do not induce tolerant growth behaviors, which would reduce the unintended consequences of nanostructured materials in medical devices, cosmetic, and industrial applications of nanoparticles. Some of the nanoparticles that did exhibit tolerant growth behaviors were liposomal nanoparticles encapsulating existing drugs or nanoparticles, and recent reports have shown that metabolic inputs may eliminate tolerant behavior. It may be possible to encapsulate metabolic inputs in liposomal nanoparticles extending the useful lifetime of the carrier and antibiotic. A rapid and accurate description of bacteria-nanoparticle interaction could be achieved by expanding our parameter extraction and statistical classification method on cell viability data using recently reported DLS methods to track cell division.

## Data Availability

To encourage further machine learning applications, all the data is available.

## Author Statement

AJ developed and implemented the data analysis and wrote the paper; DMC developed and performed experiments with bacteria derived metallic nanoparticles expect as noted below; GM developed and performed experiments with amphiphilic peptides; LBP developed and performed experiments with *H. pylori* derived metallic nanoparticles; NB developed and performed experiments with MRSA and the polymersomes; CGB and NYK developed and performed experiments with MRSA and liposomal NPs; TW proposed the analysis, advised on data collection and analysis. All authors have reviewed and revised the paper for accuracy.

## Supplementary Information

**SI Figure 1.**
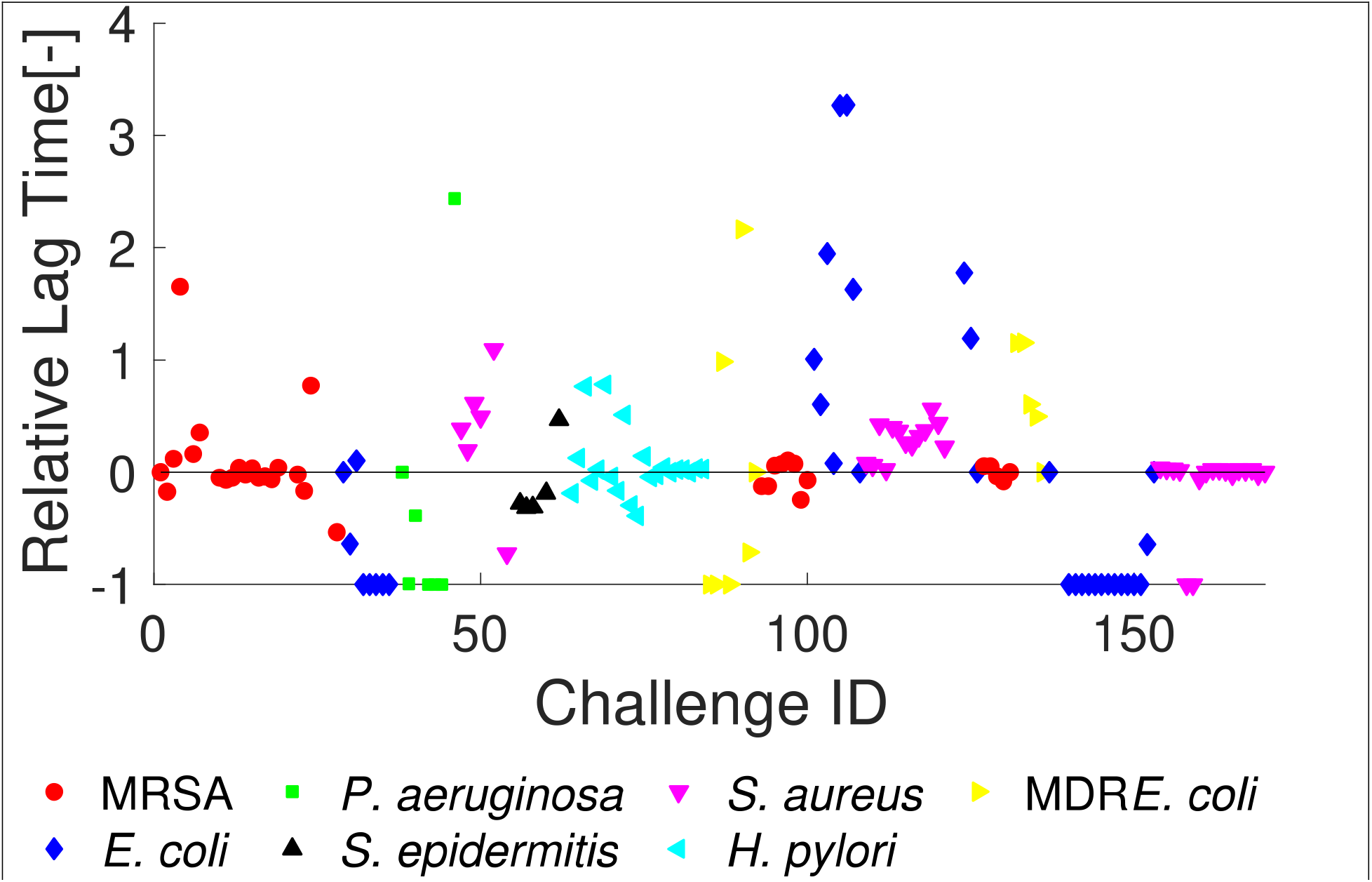
The difference in lag time relative to the control exposure,*λ* − *λ*_0_/*λ*_0_, is one of the standard methods to quantify antibiotic effectiveness.

**SI Figure 2.**
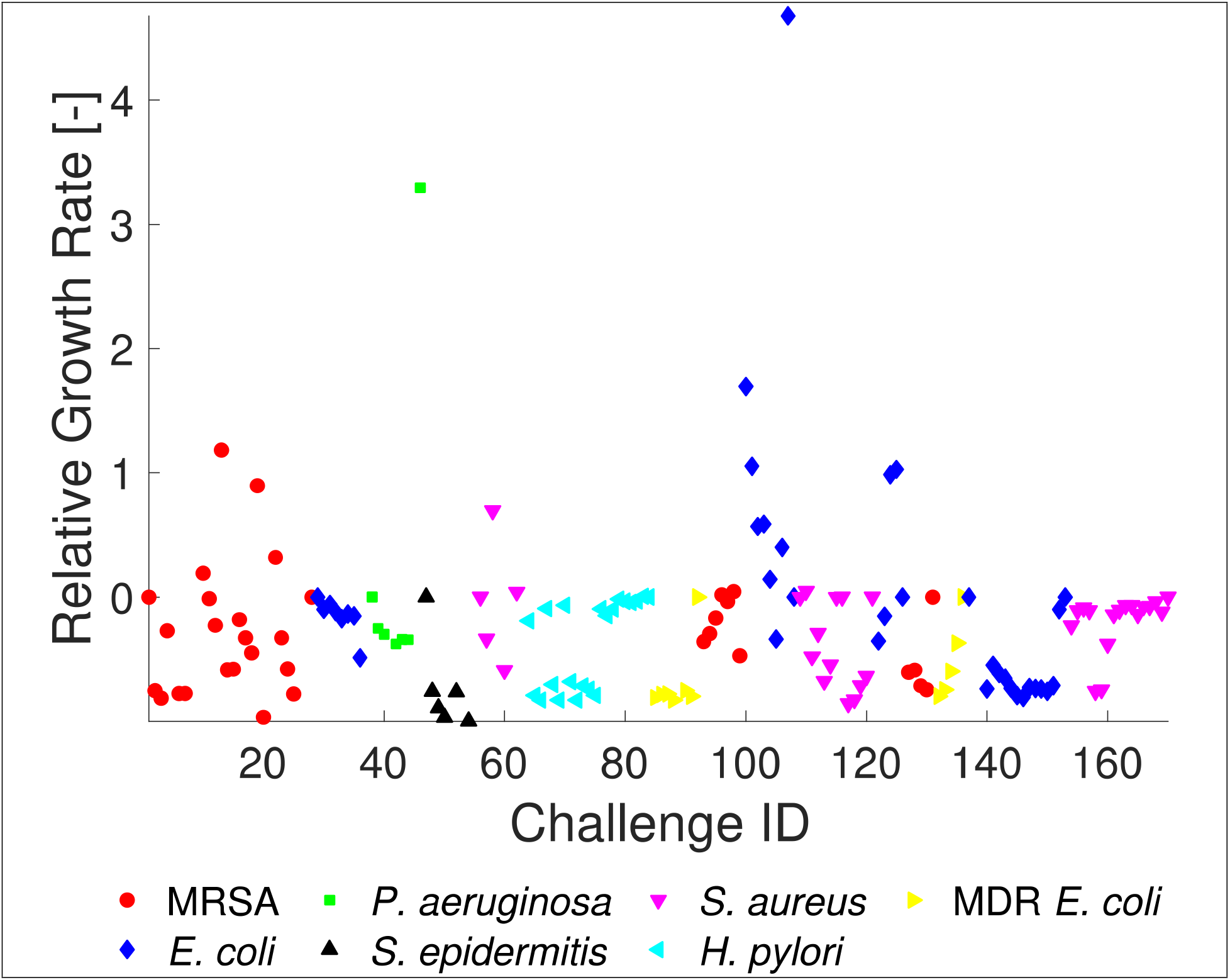
The difference in growth rate relative to the control exposure, *μ* − *μ*_0_/*μ*_0_.

**SI Figure 3.**
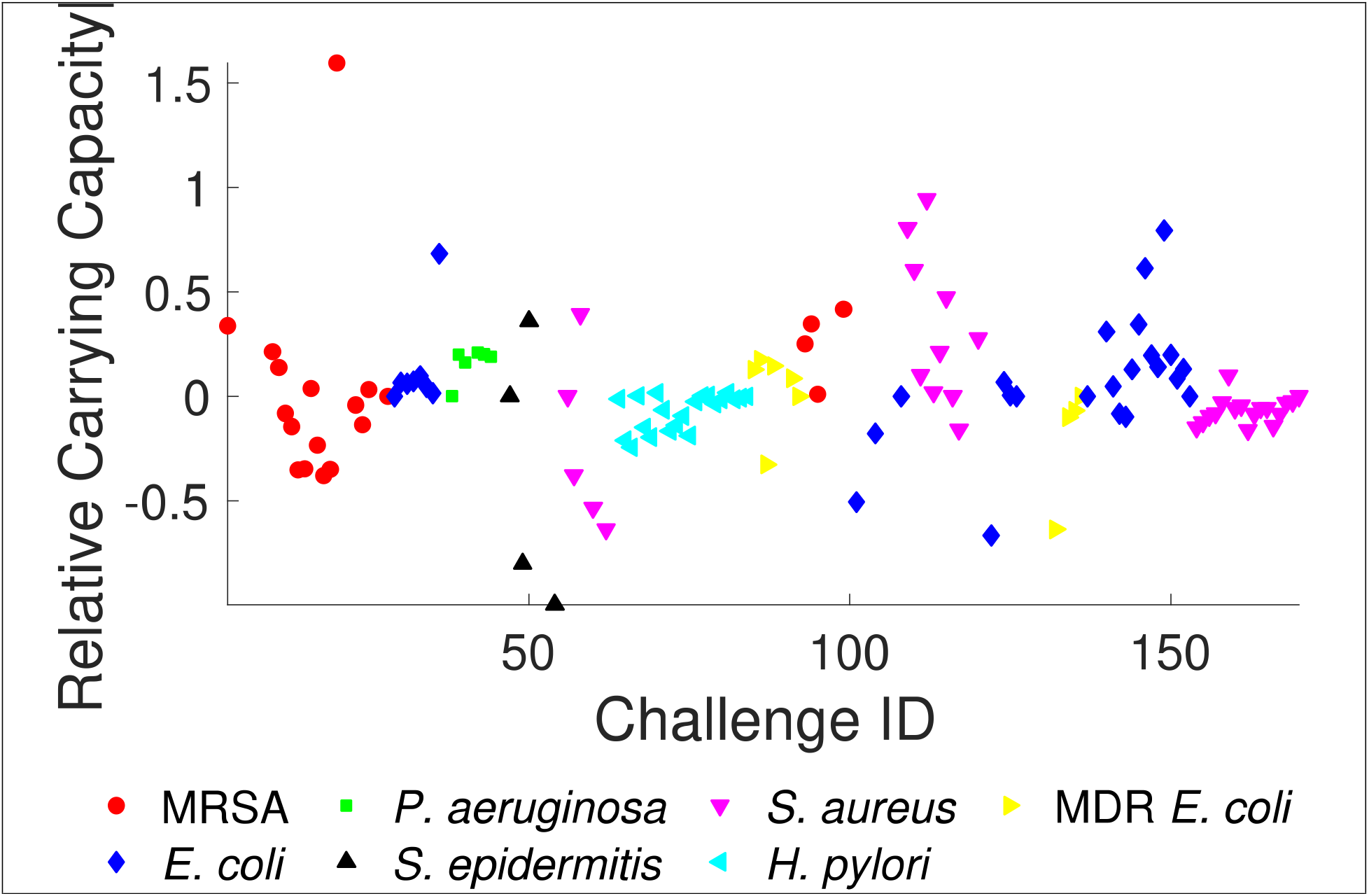
The difference in carrying capacity relative to the control exposure, *A* − *A*_0_/*A*_0_.

**SI Figure 4.**
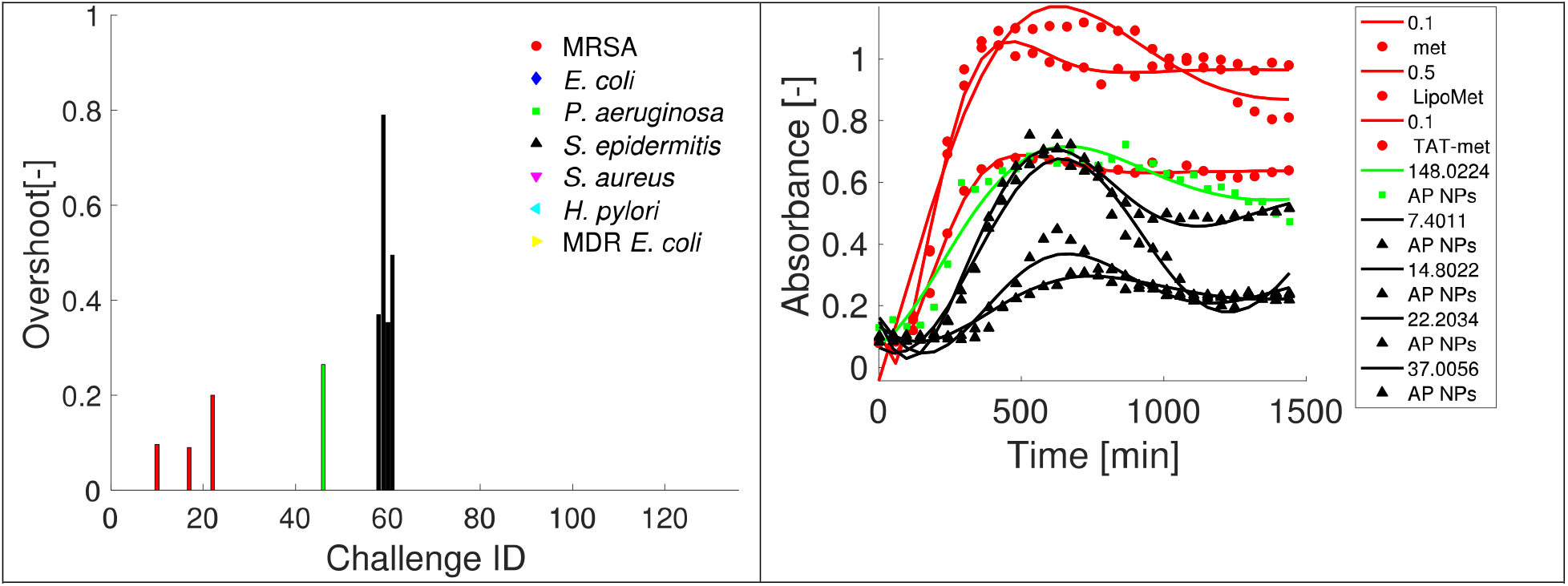
The population overshoot of bacteria and the fits of the growth curves to the model. This fit was not accepted into the model unless the AIC was the minimum of all models. This choice may have eliminated some response curves that showed oscillation but no overshoot. SI Table 1, EID # 10, 17, 22, 46, 58, 59, 60, 61.

**SI Figure 5.**
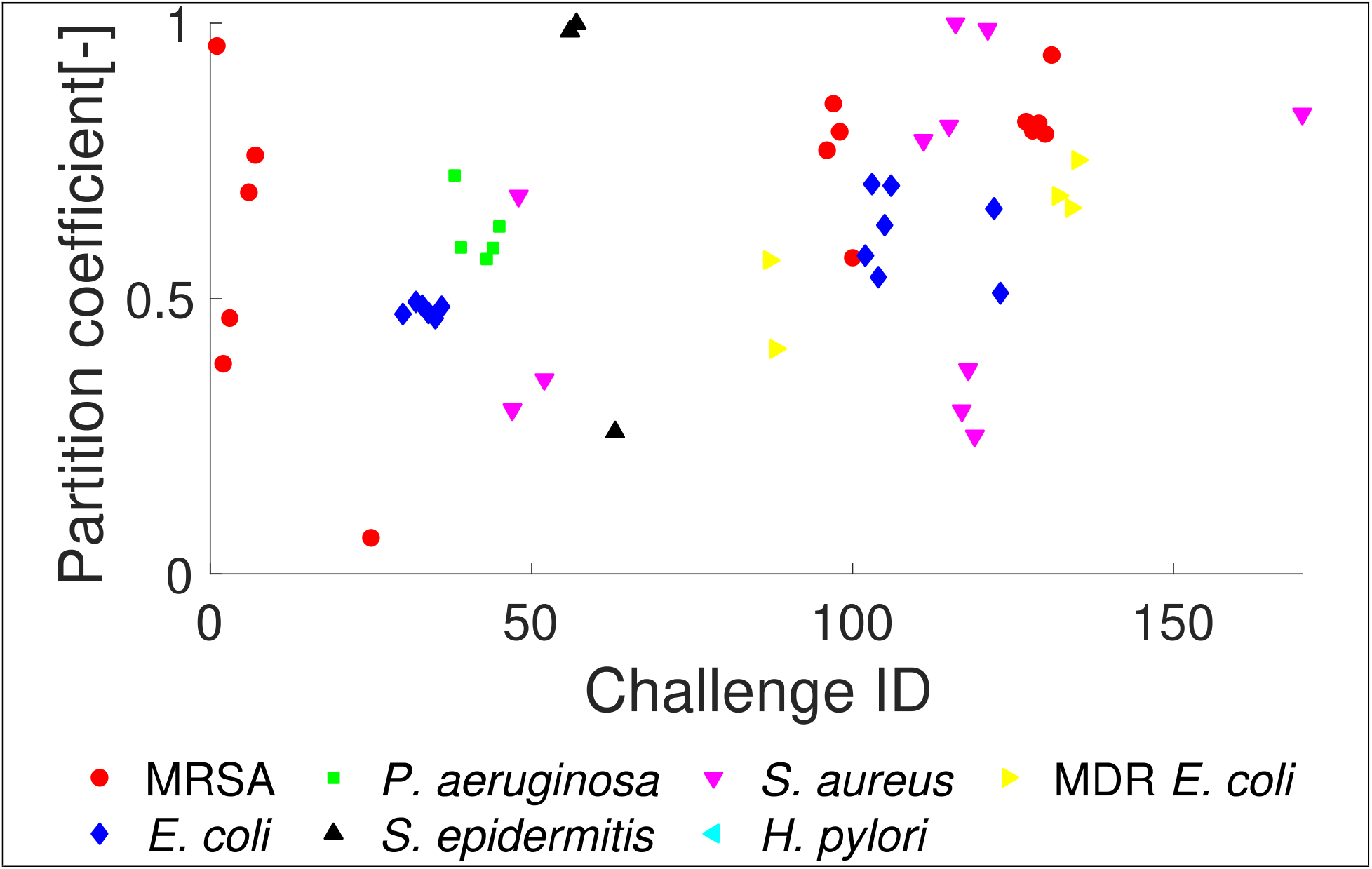
The partition coefficient describes growth curves with 0 two-phase growth, which resembles logistic growth, to 1 secondary growth phase which also resembles logistic growth. As the seven-parameter model could produce acceptable fits, the model was constrained to those where 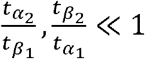 in Equation 4.

**SI Figure 6.**
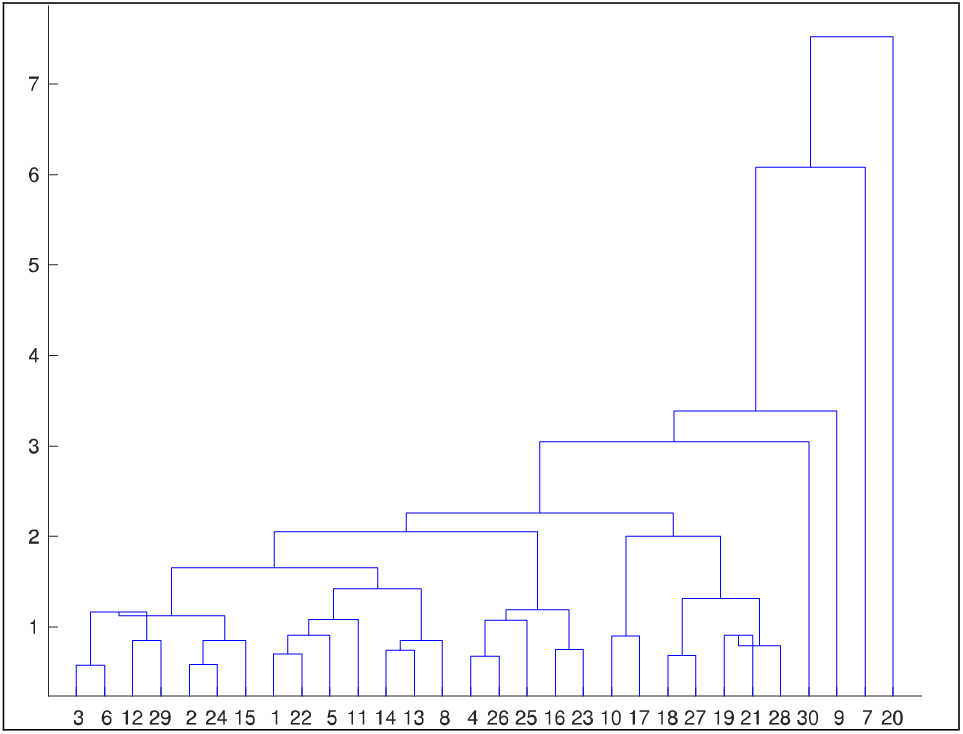
Dendogram of the hierarchical agglomerative clustering showing that the centroid clustering does not produce a monotonic cluster tree but distinct clusters, as required.

| **EID** | **Bacteria** | **NP** | **Concentrations**<br>**(ug/mL)** |
| --- | --- | --- | --- |
| 1 | MRSA | Control |  |
| 2 | MRSA | Polymersomes (PDLLA-PEG-COOH) | 1000 |
| 3 | MRSA | Polymersomes (PDLLA-PEG-COOH)<br>(methicillin) | 1000 |
| 4 | MRSA | Polymersomes (PDLLA-PEG-COOH)<br>(Ag) | 1000 |
| 5 | MRSA | Polymersomes (PDLLA-PEG-COOH)<br>(methicillin + Ag) | 1000 |
| 6 | MRSA | Polymersomes (PDLLA-PEG-COOH)<br>+PR Arginine | 1000 |
| 7 | MRSA | Polymersomes (PDLLA-PEG-COOH)<br>+PR Arginine (methicillin) | 1000 |
| 8 | MRSA | Polymersomes (PDLLA-PEG-COOH)<br>+PR Arginine (Ag) | 1000 |
| 9 | MRSA | Polymersomes (PDLLA-PEG-COOH)<br>+PR Arginine (methicillin + Ag) | 1000 |
| 10-15 | MRSA | Methicillin | 0.1, 0.5, 0.9, 1.7,<br>3.3, 5.0 |
| 16-21 | MRSA | Liposomes (methicillin) | 0.1, 0.5, 0.9, 1.7,<br>3.3, 5.0 |
| 22-27 | MRSA | Liposomes + TAT (methicillin) | 0.1, 0.5, 0.9, 1.7,<br>3.3, 5.0 |
| 28 | MRSA | Control |  |
| 29 | *E. coli* | Control |  |
| 30-37 | *E. coli* | Amphiphilic peptide | 3.70, 7.40, 11.10,<br>14.80, 22.20,<br>37.01, 74.01,<br>148.02 |
| 38 | *P. aeruginosa* | Control |  |
| 39-46 | *P. aeruginosa* | Amphiphilic peptide | 3.70, 7.40, 11.10,<br>14.80, 22.20,<br>37.01, 74.01,<br>148.02 |
| 47 | *S. epidermidis* | Control |  |
| 48-55 | *S. epidermidis* | Amphiphilic peptide | 3.70, 7.40, 14.80,<br>22.20, 37.01,<br>74.01, 148.02 |
| 56 | *S. aureus* | Control |  |
| 57-63 | *S. aureus* | Amphiphilic peptide | 3.70, 7.40, 11.10,<br>14.80, 22.20,<br>37.01, 74.01,<br>148.02 |
| 64,67,70,7<br>376,79 | *H. pylori* | H pylori produced SeNPs from 1 mM<br>Na<sub>2</sub>SeO<sub>3</sub> | 5 10, 25, 50, 75,<br>100 |
| 65,68,71,7<br>4,77,80 | *H. pylori* | H pylori produced SeNPs from 2 mM<br>Na<sub>2</sub>SeO<sub>3</sub> | 5 10, 25, 50, 75,<br>100 |
| 66,69,72,7<br>5,78,81 | *H. pylori* | H pylori produced SeNPs from 5 mM<br>Na<sub>2</sub>SeO<sub>3</sub> | 5 10, 25, 50, 75,<br>100 |
| 82-84 | *H. pylori* | Control |  |
| 85-86 | MDR *E. coli* | Liposomes (cysteine capped AgNPs) | 5, 25 |
| 87-88 | MDR *E. coli* | Liposomes (Glutathione capped AgNPs) | 5, 25 |
| 89-91 | MDR *E. coli* | Cysteine capped AgNPs | 5, 25, 50 |
| 92 | MDR *E. coli* | Control |  |
| 93-94 | MRSA | Liposomes (cysteine capped AgNPs) | 5, 25 |
| 95-96 | MRSA | Liposomes (Glutathione capped AgNPs) | 5, 25 |
| 97-99 | MRSA | Cysteine capped AgNPs | 5, 25, 50 |
| 100 | MRSA | Control |  |
| 101-102 | *E. coli* | Liposomes (cysteine capped AgNPs) | 5, 25 |
| 103-104 | *E. coli* | Liposomes (glutathione capped AgNPs) | 5, 25 |
| 105-107 | *E. coli* | Cysteine capped AgNPs | 5, 25, 50 |
| 108 | *E. coli* | Control |  |
| 109-110 | *S. aureus* | Liposomes (cysteine capped AgNPs) | 5, 25 |
| 111-112 | *S. aureus* | Liposomes (glutathione capped AgNPs) | 5, 25 |
| 113-115 | *S. aureus* | Cysteine capped AgNPs | 5, 25, 50 |
| 116 | *S. aureus* | Control |  |
| 117-120 | *S. aureus* | Glutathione capped AgNPs | 6.9, 13.8, 69, 138 |
| 121 | *S. aureus* | Control |  |
| 122-125 | *E. coli* | Glutathione capped AgNPs | 6.9, 13.8, 69, 138 |
| 126 | *E. coli* | Control |  |
| 127-130 | MRSA | Glutathione capped AgNPs | 6.9, 13.8, 69, 138 |
| 131 | MRSA | Control |  |
| 132-135 | MDR *E. coli* | Glutathione capped AgNPs | 6.9, 13.8, 69, 138 |
| 136 | MDR *E. coli* | Control |  |
| 137 - 140 | *E. coli* | MRSA-SeNPs | 250,150,75,25 |
| 141 - 144 | *E. coli* | *S. aureus*-SeNPs | 250,150,75,25 |
| 145 - 148 | *E. coli* | *E. coli*-SeNPs | 250,150,75,25 |
| 149 - 152 | *E. coli* | *P. aeruginosa*-SeNPs | 250,150,75,25 |
| 153 | *E. coli* | Control |  |
| 154-157 | *S. aureus* | MRSA-SeNPs | 250,150,75,25 |
| 158-161 | *S. aureus* | *S. aureus*-SeNPs | 250,150,75,25 |
| 162-165 | *S. aureus* | *E. coli*-SeNPs | 250,150,75,25 |
| 166-169 | *S. aureus* | *P. aeruginosa*-SeNPs | 250,150,75,25 |
| 170 | *S. aureus* | Control |  |

